# Emphasizing the ‘positive’ in positive reinforcement: Using non-binary rewarding for training monkeys on cognitive tasks

**DOI:** 10.1101/169201

**Authors:** Benjamin Fischer, Detlef Wegener

## Abstract

Non-human primates constitute an indispensable model system for studying higher brain functions at the neurophysiological level. They can be trained on highly demanding cognitive tasks, and studies involving these animals elucidated the neuronal mechanisms of various cognitive and executive functions, such as visual attention, working memory, and decision-making. The training of behavioral tasks used to study these processes builds on reinforcement learning and involves many discrete stages. It may takes several months, but frequently lasts a year or longer. The training is usually based on applying a liquid reward as the reinforcer to strengthen the desired behavior, and absence of the reward if the animal’s response was wrong. We here propose an alternative, non-binary rewarding scheme that aims to minimize unrewarded behavior. We show the potential of this alternative scheme to significantly speed up the training of an animal at various stages, without trade-off in accessible task difficulty or task performance.

## Introduction

Non-human primates are capable to learn sophisticated behavioral tasks. During the last decades and since, insights into the neuronal mechanisms of cognitive control and executive functions were gained by physiological experiments requiring animals to perform specific cognitive operations while neuronal activity was recorded in some cortical area of interest. The behavioral tasks used in such studies are often complex and highly demanding. They frequently require to follow intricate selection and response rules (Herrington and Assad 2009; Schledde et al. 2017; Shichinohe et al. 2009; Watanabe and Sakagami 2007), may and include storage of numerous items in working memory (Barone and Joseph 1989; Inoue and Mikami 2006; Miller et al. 1996), sustained covert attention to peripheral stimuli (Hafed et al. 2011; Wegener et al. 2004), multiple-object tracking in space (Matsushima and Tanaka 2014; Mitchell et al. 2007b), identification of a target shape in a morphing object (Grothe et al. 2012; Taylor et al. 2005), and even video gambling with another monkey (Hosokawa and Watanabe 2012) and basic arithmetic (Cantlon and Brannon 2007). Training of such tasks (and, likewise, the training of much simpler forms of cognitive tasks) is usually acquired by a succession of many small steps, with each step requiring adaptation of the learned behavior to a slightly changed task rule (successive approximation).

Positive reinforcement training (PRT) constitutes the gold standard for animal training procedures. A substantial body of literature is available describing the benefits of PRT for scientific, veterinary, and husbandry purposes (for a tabulated overview, see Prescott et al. 2005), but only a few papers deal with the specifics of training procedures for cognitive tasks in the laboratory (Mitchell et al. 2007a; Remington et al. 2012; Scott et al. 2003; Watanabe and Funahashi 2015). The usual procedure, however, consists of reward delivery in response to the desired behavior (e.g. a fast lever release following the onset of a target stimulus), and withholding of reward for the wrong behavior (e.g. a lever release that comes too soon or too late), typically accompanied by trial termination (Newsome and Stein-Aviles 1999). We will call this form of feedback a ‘binary’ reward schedule in the remainder of the article, since it consists of either delivering or not delivering the reinforcer to the animal. We will argue that such binary rewarding might be sub-optimal for quick and effective learning under laboratory conditions.

Among the many factors influencing the overall time needed to train an animal on a demanding cognitive task, three are outstanding: i) the overall motivation of the animal, ii) the effective number of trials the monkey can learn on, and iii) the unambiguousness of the feedback. A pure binary reward scheme has obvious disadvantages with respect to each of these. First, regarding motivation, consider that monkeys have no insight into the structure and the requirement of a new task (or task rule), and depend on learning by trial-and-error. Hence, many trials will terminate with the undesired response and the monkey will not be rewarded. Since the animal may have spent some effort to perform the trial correctly, withholding a reward potentially counteracts the intrinsic motivation to perform the task. Second, despite this negative influence of errors, they constitute the most important source of information during learning. Only if a trial does not end with the expected reward, there is a need for the animal to adapt its behavior. Yet, high error rates bear the risk that the animal will end a training session early, and/or gets used to errors on the long-term. In turn, small error rates will reduce the amount of information the monkey can build on for adapting to a new task rule. For quick and effective learning, none of these alternatives is optimal. Third, withholding the reward and termination of a trial in response to an error constitutes an ambiguous feedback signal: it is signaling the animal that its response was not the desired one, but provides no hint for the source of the error.

We here propose an alternative, non-binary reward scheme that emphasizes successful behavior (NB-PRT, non-binary positive reinforcement training). For any new training step, the to-be-learned behavior gets associated with a high amount of reward, while other behavior may receive lesser, graded reward. We argue that this reward regime helps to refine training procedures and serves the 3R principle (Russell and Burch 1959) by i) providing a more differentiated feedback, ii) increasing the efficient number of trials the monkey can learn on, iii) reducing the number of unrewarded trials, and iv) providing more information for the trainer to find the best suited training schedule. The paper explains the details of this approach, and provides four examples for its efficiency, including situations where it may seem to be counterproductive at first view. The paper does not attempt to provide a quantitative comparison of the two different reward schemes outlined above, but rather to serve the community with the experiences gathered with this alternative approach.

## Material and Methods

The training procedures described here were carried out with two male macaque monkeys (Macaca mulatta) of about 10 and 14 kg body weight, implanted with a headpost. All surgical and experimental procedures followed the Regulation for the Welfare of Experimental Animals issued by the Federal Government of Germany, and were approved by the local authorities. Surgical procedures followed previously published protocols (Schledde et al. 2017; Wegener et al. 2004). Out of training sessions, animals were kept in a species-appropriate, environmentally enriched husbandry. They received free fruits and water during the weekend and during non-training periods.

Training sessions were performed in a lightly dimmed room. Animals sat in a primate chair 80 cm in front of a 22-inch CRT display (1280 x 1024 pixels, 100 Hz refresh rate). Eye movements were measured at a spatial resolution of 0.2 deg visual angle, using a custom-made remote video-oculography system. Reward consisted of water or diluted red grape juice, applied by a simple, valve-controlled gravity liquid dispenser. A “high reward” consisted of about 25 – 30 ml/100 correct responses, a “medium reward” of about 15 – 20 ml/100 corrects, a “small reward” of about 10 – 15 ml/100 corrects, and a “very small reward” of about 5 – 10 ml/100 corrects.

Visual stimulation was carried out with custom-made software, run on a Pentium computer. The computer software chose the amount of reward depending on the monkeys’ behavior (e.g. precision of fixation or reaction time), and saved all stimulation and behavioral data in a trial-description file. Data were analyzed using Matlab (The Mathworks, Natick, MA). Training sessions with less than 50 trials were disregarded. For better traceability, specific task requirements are explained in the corresponding Results sections.

Data analysis was based on the amount of correct and error responses, and reaction time (RT) distributions. RT distributions were fit by an ex-Gaussian probability density function, which is a convolution of a Gaussian and an exponential distribution. The ex-Gaussian fit provides the parameters μ and σ to describe the mean and the variability, respectively, of the normally distributed component of the distribution, and parameter τ to describe the exponential part, which accounts for the skew of the distribution (Heathcote et al. 1991; Tarantino et al. 2013).

Statistical testing was done using unpaired, two-sided *T*-tests. RT analysis was based on the medians of the respective sessions.

Regarding the procedure described with reference to Figure 4, changes at cued objects were delayed by at least 750 ms from the preceding change at the uncued object. Trials in which such uncued changes occurred are referred to as catch trials, and were kept at a rate of 10% to 20%. Responses to both cued and uncued changes were allowed during a 550 ms time window starting 200 ms after the change. For some sessions at the beginning of the training to introduce catch trials, the minimal temporal separation between changes at uncued and cued objects was allowed to be as short as 450 ms. In such cases, when the temporal separation between two changes was too short to allow for a clear assignment of the response, we considered responses below 750 ms after the first change as a response to the first, uncued change, and responses occurring later as a response to the second, cued change. Sessions with less than 25 catch trials were not considered for catch trial analysis. For the analysis of reward amount and RT in catch trials, we normalized the data to their corresponding median value from normal trials (trials without catch event) of the same session. Thus, a value of 1 indicates that the reward amount or RT was the same as in normal trials, and values below or above 1 indicate that it was smaller or larger, respectively, than in normal trials.

## Results

In the following, we provide four examples for non-binary rewarding during training of different tasks/task rules. Based on studies of reward-based decision making (Amiez et al. 2006; Feng et al. 2009), our approach builds on the assumption that monkeys will effectively adapt their behavior to maximize the reward amount delivered in a single trial, and across a series of trials. The first example will consider a relatively simple requirement for precise fixation during covert attention, while later examples will deal with shaping selective attention during progressively more complex task conditions. In each example, we show that NB-PRT can exert positive effects regarding the number of trials the monkey can learn on, the quality of the feedback, the overall motivation of the animal to participate in the training, and its confidence with the task’s requirements.

### Example 1: Improving fixation behavior

Consider a monkey having learned to properly gaze at a central fixation point (FP) and press a lever to initiate a trial, keep its gaze at the FP, and release the lever in response to the dimming of the FP after a pseudo-random interval of up to several seconds. Now assume this monkey shall be trained to attend a peripheral target while keeping proper fixation, and to detect a sudden feature change of this target. Compared to what was learned previously, this task has two new components that are likely to cause errors: first, keeping fixation while covertly attending to a distant location, and second, detect a feature change of a distant stimulus instead of the FP dimming. Such a combination of unknown error types is a critical step in the training, particularly for unexperienced animals during early training steps. If errors become too frequent, monkeys may get frustrated and perform fewer trials than before, or get used to errors and only slowly learn the desired behavior.

Usually, we train such a new response criterion by placing the target close to the FP and let a salient feature change occur 200 ms before the FP dimming. This makes the change become an easy-to-detect predictor of the more difficult-to-detect FP dimming, and the monkey will quickly use this information to initiate its response. Online analysis of the RT distribution allows monitoring the learning progress of the monkey, and a leftward shift of the RT distribution indicates that the monkey starts to respond to the predictor. The target’s feature change would then replace FP dimming. This can usually be achieved within a single training session. The more critical step is to move the target into the periphery, and to place it at various locations. The requirement of covertly attending a peripheral target results in a (strongly) increased likelihood of fixation errors. Using a binary reward scheme, errors caused by imprecise fixation or saccading towards the target are not differentiated from errors related to the new task rule. For the animal, the combined effect of ambiguous feedback (due to multiple possible (new) error causes) and more unrewarded trials is likely to induce a reduction in confidence with the current task rules and overall motivation, respectively. Preventing this requires to split the training into small steps, e.g. by slowly approaching the final target position over the course of several training sessions. This is likely to work but time-consuming.

NB-PRT provides more opportunities. Instead of dividing the training into small steps, we divide the reward into different amounts. For the example outlined above, we trained the new task rule (detect a speed-change of an inherently moving Gabor grating) during one session, and then placed the Gabor at its desired final location in the first session following this training (Fig. 1*A*). To prevent ambiguous feedback and to keep the rate of successful trials high, we divided the fixation window into two zones: an inner fixation window with a radius of 1 deg (as already used in previous training), and an outer fixation window with a radius of 2 deg. For precise fixation, the monkey obtained a medium reward, and for unprecise fixation, it obtained a small reward (see Material and Methods for nomenclature of rewards and associated amounts). We continued to treat saccades of more than 2 deg amplitude (leaving the outer fixation window) as errors, resulting in the termination of the trial. In the first and second training session, the monkey made about 25% fixation errors (Fig. 1*B*), due to saccading towards the target stimulus. Based on reports from human subjects performing this task (Wegener et al. 2008), we considered the monkey being aware of such large saccades and the error signal provided a clear feedback. In contrast, small fixation errors during initial training of a covert attention task may occur without the animal being aware of these. Not terminating but instead rewarding these trials with a reduced amount of liquid prevented an unnecessarily high error rate, kept the motivation of the animal to perform the task, and still provided the monkey with information about the desired behavior due to graded rewarding of successful trials as a function of fixation precision.

**Figure 1.**
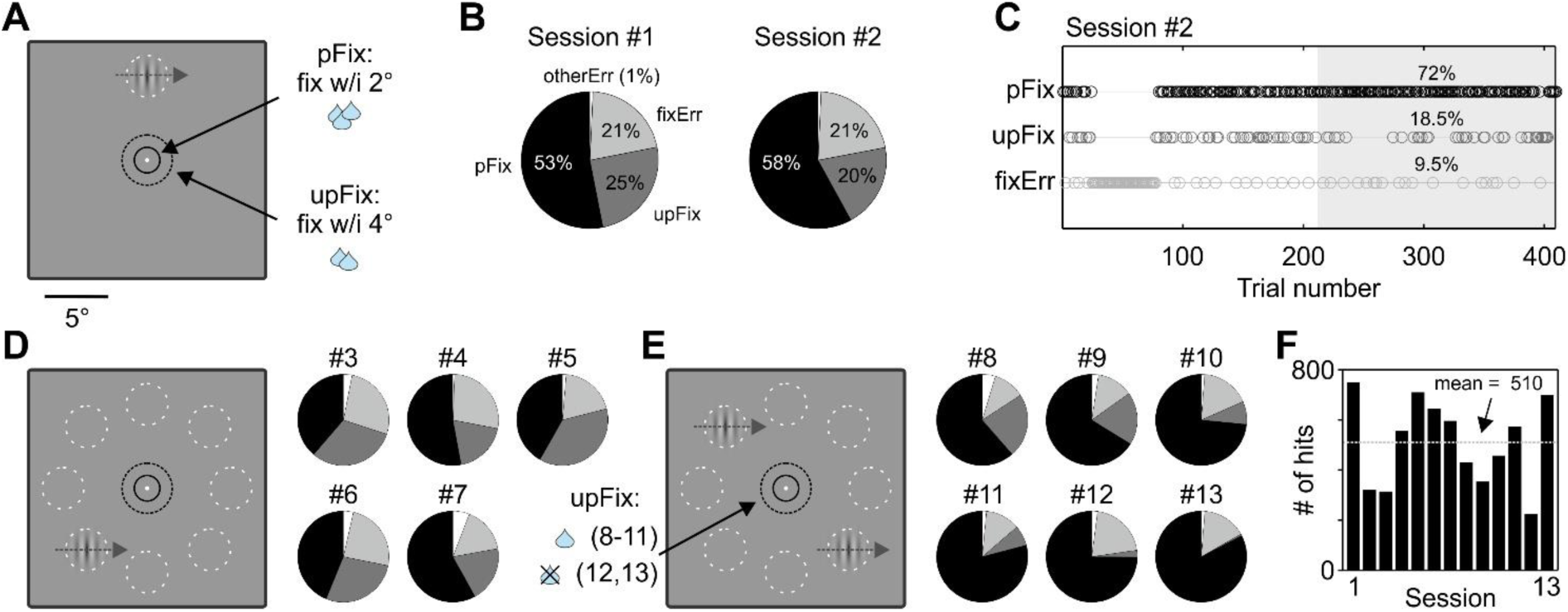
Non-binary reward scheme during training of a covert attention task. (*A*) Stimulus display, fixation windows, and reward associations (pFix: precise fixation; upFix: unprecise fixation) during training sessions 1 and 2 after the animal had learned to release the lever in response to the speed-change of a Gabor stimulus with inherent motion. (*B*) Mean behavioral performance (fixErr: fixation errors; otherErr: errors not related to fixation (false alarms and misses)). (*C*) Behavioral performance during session 2. Percentage values provide performance during the last 200 trials (gray-shaded area). (*D* and *E*) Visual display and possible target positions during sessions 3 to 7 (*D*), and 8 to 13 (*E*), and behavioral performance. Pie shading corresponds to (*B*). (*F*) Number of correct responses during all training sessions. Dashed horizontal line indicates their mean. Throughout the figure, solid black circles in display diagrams indicate ‘precise fixation zone’ (pFix, 2 deg diameter); dashed black circles indicate ‘unprecise fixation zone’ (upFix, 4 deg diameter); dashed grey circles indicate possible target positions. Circles are shown for illustration purposes only, they were not displayed. Drop-symbols: 3 drops, medium reward; 2 drops, small reward; 1 drop, very small reward.

During the beginning of session 2, the monkey made a long series of eye errors, presumably trying a different behavioral strategy to get the maximal reward (Fig. 1*C*). Yet, assuming that we chose the size of the outer fixation window properly, and the monkey was indeed capable to distinguish trials with a saccade towards the target from trials in which it largely kept fixation, the feedback signal kept clear and in line with the previously learned rule of gazing at the FP. As such, the monkey went back to the previous strategy, reduced the number of errors instantaneously, and then adapted its behavior in the desired manner. During the last 200 trials of the same session, the monkey performed above 70% of all trials with precise fixation, and less than 10% of the trials were terminated due to an eye error (Fig 1*C*). Thus, the monkey shaped its behavior in the intended manner within two sessions, even though attending covertly to a peripheral object and responding to a speed change constituted new task requirements.

With this result, we went on to the next training step and placed the target at eight different positions, each at a distance of 8 deg from the FP. We also introduced a cue to indicate the upcoming target position at the beginning of the trial (low contrast first and gradually increased during session 3). We trained these positions for five sessions during which the monkey constantly made only a small number of errors not related to fixation, and a moderate number of fixation errors (16 – 27%). The ratio of precise fixation trials among all successful trials was between 53% and 75% (Fig. 1*D*). During sessions 8 to 13, we additionally presented a second stimulus opposite to the target (low contrast first and gradually increased during session 8), and reduced the amount of reward for unprecise fixation to “very small” in sessions 8 to 11. In session 12 and 13, such trials were not rewarded anymore, but neither were they terminated. At this stage, we considered the desired behavior as fairly well established, and withholding of reward for trials with unprecise fixation as not likely to induce uncertainty about the task rules. Not rewarding unprecise fixation trials instead more strongly forced the monkey to fixate precisely. In line with this, during sessions 12 and 13 the number of hits with precise fixation reached 75% and 83% of all trials, and the number of eye errors did not exceed 21% and 15%, respectively (Fig. 1*E*).

Thus, within 13 training sessions the monkey learned to covertly attend to one of several peripheral locations with good fixation performance, and to reliably indicate a speed-change of a peripheral target stimulus in the presence of a distracting second stimulus. The non-binary reward regime allowed the monkey to figure out on its own how to get the highest reward, and constrained errors to previously established rules (here: suppression of large saccades). In our past attempts, using binary rewarding was much slower to train this behavior. Limiting the fixation window to promote precise fixation usually caused a strong increase in the error rate and an accompanying reduction of the animal’s confidence with the task requirements, while increasing the fixation window did not support precise fixation. NB-PRT, in contrast, kept the animal’s motivation high and resulted in much quicker learning and an average number of more than 500 successful trials during this training block (Fig. 1*F*), even though single trials lasted for up to 5 sec.

### Example 2: Focusing attention

Training progress critically depends on the effective number of trials the monkey can learn on. NB-PRT can be used to increase their efficient ratio (i.e. independent from the absolute number of trials the monkey is performing). In our example, we continue the training situation described in the previous section (Fig. 1*E*). The next step was to increase the task’s attentional demand. During sessions 12 and 13, when the monkey was already good in performing the task in the desired manner, we switched back to binary rewarding to reset the reward regime before the next training step. During these sessions, the monkey obtained a fixed, medium reward for detecting the target’s speed change within 150 to 750 ms without leaving the inner fixation window. It reached a performance of 81% correct responses and a median reaction time (RT) of 380 ms (Fig. 2*A*). To force the monkey to strongly focus its attention on the target object, it shall learn next to respond as fast as possible. With a binary reward scheme, this might be achieved by reducing the length of the response time window. Consequently, a fraction of previously successful responses will become future errors, and these errors provide the information for the monkey to adapt its behavior. In the example of Figure 2*A*, reducing the maximal response time from 750 to 450 ms would label about 9% of the trials as misses. Even if assuming the animal will understand the cause of these errors (and not confuse them with errors of different source), it must perform many trials to arrive at one “teacher trial”. Increasing the number of teacher trials for faster learning is obligatorily linked to a higher number of errors, and again likely counteracting the confidence and the overall motivation of the animal if errors exceed a critical number. Moreover, with this schedule the monkey learns to not be slow, which is obviously not the same as being fast.

**Figure 2.**
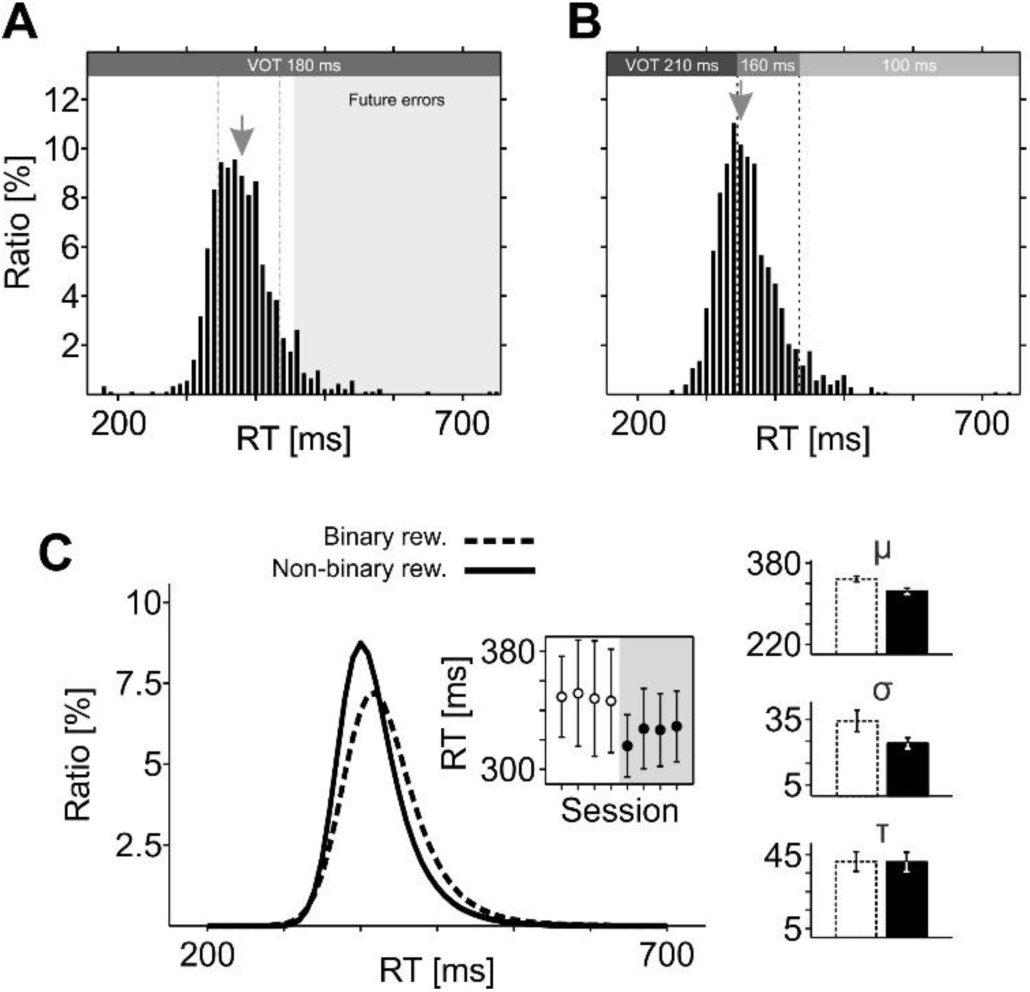
RT-dependent rewarding. (*A*) Empirical RT distribution for speed-change detection with binary reward, averaged over 2 days before introducing RT-dependent rewarding. Gray rectangle indicates future errors if limiting the response window to 450 ms. Dashed gray lines indicate response time windows as shown in (*B*). VOT, valve opening time (reward amount); gray arrow, median RT. (*B*) Empirical RT distribution averaged over two days following a one-session introduction of RT-dependent rewarding. Grey-shaded horizontal bars indicate VOT for fast, medium, and slow RTs. (*C*) Ex-Gaussian fits of RT distributions from four sessions prior to introducing RT-dependent rewarding, and four sessions thereafter. Inlet shows the μ and σ-values of single sessions. Bar plots on the right show the mean μ and σ-values of the Gaussian component, and the mean τ of the exponential component before (left) and after (right) introducing RT-dependent rewarding. Errors bars, S. D.

NB-PRT allows emphasizing the positive, to-be-facilitated behavior - the fast response. Rather than relying on the failure to respond in time, providing a large reward for every fast response results in a considerably higher number of teacher trials. For example, 25% of all trials shown in Figure 2*A* had an RT of less than 350 ms. To speed up response times, we associated such trials with a high reward, while trials with an RT of more than 440 ms received a small reward. Trials with an RT in-between were slightly less rewarded as compared to previous training sessions, to make the reward difference between fast and medium RTs clearly distinguishable. This new reward regime was highly effective and caused a decrease in the median RT from 380 ms to 350 within a single training session. Figure 2*B* shows the RT distribution of the following two sessions, revealing a total of 45% successful responses being faster than 350 ms. Ex-Gaussian fits to the four sessions before and after introducing the RT-dependent reward illustrate the clear, leftward shift of the RT distribution (Fig. 2*C*). The mean of the Gaussian component (fitting the left (fast) part of the skewed RT distribution) decreased from 348 ± 34 ms to 324 ± 24 ms (*T*-test, *P* < 10^−3^, *N* = 4), while the mean of the exponential component (fitting the right (slow) part of the skewed distribution) was unchanged (τ_binary_: 41 ms; τ_non-binary_: 41 ms; *P* = 0.9486). Thus, emphasizing the desired, fast response by RT-dependent rewarding instead of training down the undesired, slow response by reward rejection, resulted in an essentially instantaneous and lasting acceleration of response times.

### Example 3: Teaching abstract cues

Many behavioral tasks to investigate higher cognitive functions rely on abstract cues to indicate the required behavior. For example, spatial, colored, or symbolic cues may be used to indicate the identity of an object to be kept in memory (Chelazzi et al. 1998; Inoue and Mikami 2006), the type of a desired motor action (Everling and Munoz 2000; Snyder et al. 2000), or the task rule (Assad et al. 2000; Schledde et al. 2017). For the example given in the previous section, a spatial cue was used to direct the monkey’s attention to the location of one of the two simultaneously presented stimuli (Fig. 3*A*). Depending on the cue, the stimulus over the receptive field of the recorded neuron may be attended or unattended, thus allowing analysis of neuronal activity under identical physical stimulation but different loci of covert attention. Yet, to test whether the monkey follows the cue instructions and in fact selectively attends the cued item, it is necessary to introduce target events (change in color, motion direction, orientation, or the like) at uncued items, which have to be ignored by the animal. With binary rewarding, responses to such uncued events are considered as errors, resulting in the termination of the trial (Galashan et al. 2013; Reynolds et al. 1999; Treue and Maunsell 1996; Wegener et al. 2004).

**Figure 3.**
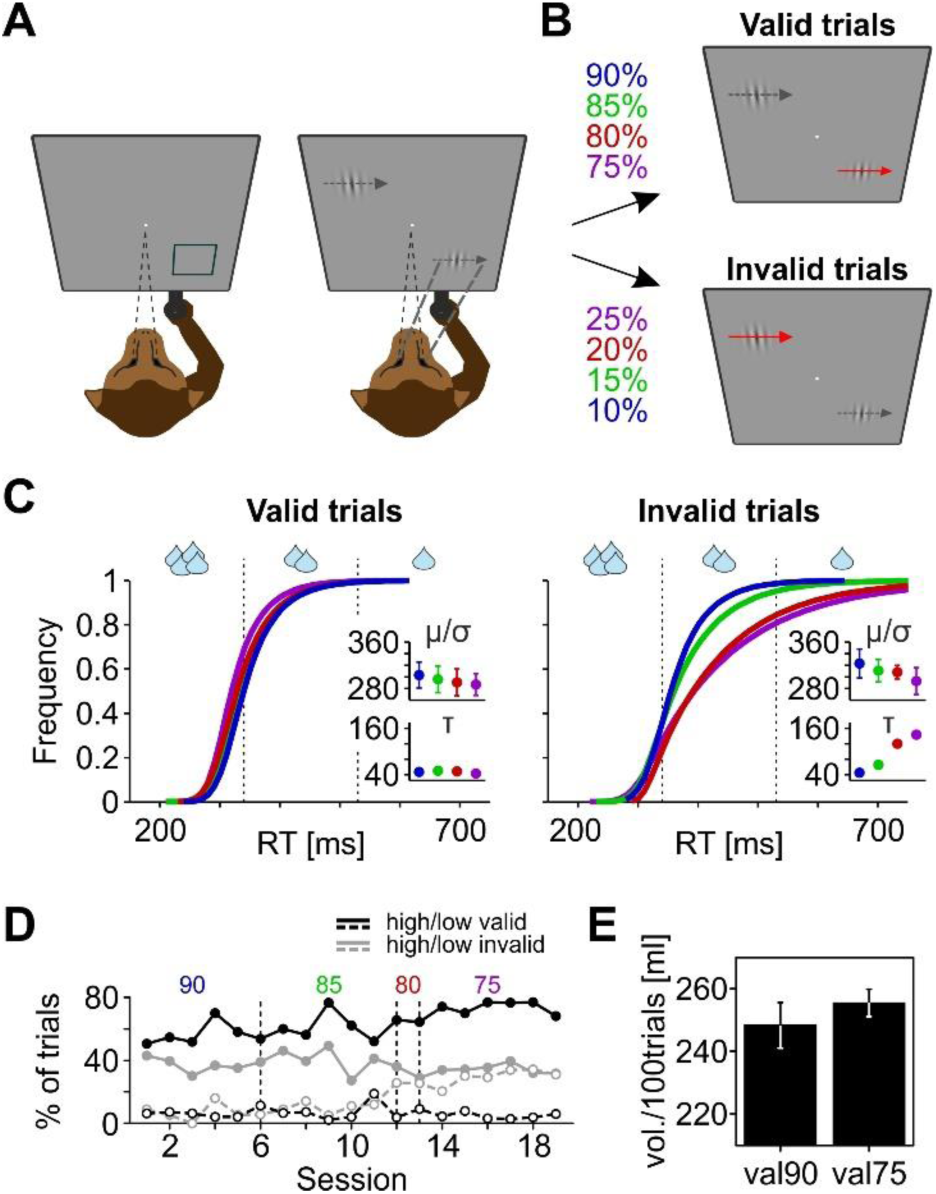
Training of abstract cues. (*A*) Visual display. The monkey was required to gaze at the FP in the center of the display. A rectangular frame indicated the mutual position of the object to undergo the to-be-detected speed-change event (left). Gabor stimuli appeared subsequently at two FP-mirrored locations (right). The desired behavior was to use the cue information for allocating spatial attention (bold dashed lines). (*B*) In valid trials, the speed change occurred at the cued location, while it occurred at the uncued object in invalid trials. Validity was changed over the course of training sessions. (*C*). Empirical RT distributions in valid and invalid trials. Inlets depict the mean parameters of Ex-Gaussian fits to each of the RT distributions, depending on cue validity. Color code as in (*B*). (*D*) Relative amount of reward obtained valid and invalid trials, depending on cue validity. Trials with medium reward are disregarded for simplicity. (*E*) Average reward amount per 100 trials during early (90% validity) and late (75% validity) training sessions. Error bars, S.D.

Training the meaning of such abstract cues frequently is a hard test of patience. From the perspective of the trainer, the task rule might be straightforward since reward is only provided for responses to the cued item. Yet, from the perspective of the animal, the situation is less clear. Even with selective attention directed to the cued item, a target event occurring at the uncued item usually is not out of sight but may be well perceived (Braun and Sagi 1990; Wegener et al. 2008). Responding to this event is exactly what has been trained before – for the example of Figure 3*A*, the monkey would perfectly be in line with the previously learned task rules if indicating a speed change of any of the two stimuli, independent of the cue. With a binary reward, there is no clear alternative to guide the animal towards the desired behavior than adjusting the number of errors by some means - hoping that at some point the animal gets the association between cue appearance at one of the stimulus locations, and reward delivery following a correct response to that stimulus. Again, this may go along with the monkey becoming either unsure about task requirements, frustrated, or used to errors. NB-PRT, in contrast, provides additional options for shaping the monkey’s behavior, relying on the monkey’s natural expertise – getting the highest reward. In the following, we provide two examples of slightly different technical implementation, showing that abstract cues can effectively been learned within a few sessions.

The first example continuous the training situation from the previous sections. The spatial cue indicating the position of the upcoming target had been introduced when adding the second stimulus to the display, as mentioned previously. Yet, the cue had no obvious meaning with respect to uncued events, since such events were not present in earlier sessions. To use non-binary rewarding for teaching the meaning of the cue, we chose a Posner paradigm-like design (Petersen et al. 1987; Posner 1980) and presented speed changes at the uncued location in a fraction of trials. We rewarded responses to such uncued events in the same way as responses to cued events, with high, medium, and small reward for fast, medium, and slow RTs, respectively. The rationale of this scheme was to test whether the monkey would distribute attention over the whole display (likely resulting in RTs of medium length and a relatively save amount of medium reward), or alternatively, use the information provided by the cue and direct attention to one stimulus selectively (increasing the chance of a high reward for fast RTs, at the risk of getting a small reward for slow RTs if changes occur outside the attentional focus). If the monkey mainly relies on maximal reward on the short term (i.e. in single trials), and/or the reward regime is chosen to provide a higher mean reward per trial on the long-term (i.e. over the entire session), choosing the second of the two options will be highly beneficial.

We started with a cue validity of 90% and then reduced it to 85%, 80%, and 75% (Fig. 3*B*). Each validity was kept for 6 sessions, with the exception of the 80% condition, which was applied in only one session. To foster selective attention, we reduced the speed change magnitude from 100% to 80%. RTs were analyzed by fitting ex-Gaussians to their distributions for different cue validities. With reduced validity, RTs to correctly cued changes became increasingly faster (μ(σ)_[90 85 80 75]_: 303 (22), 295 (23), 291 (23), 287 (19) ms; *T*-test_(val90, val75)_, *P* < 10^−3^, *N* = 6), while RTs to incorrectly cued changes became increasingly slower, mainly due to an increase in the exponential component of the distribution (τ_[90 85 80 75]_: 45, 66, 120, 143 ms; *P* < 10^−4^). During the 90% validity condition, the median RT difference between correctly and incorrectly cued changes was 20 ms, and increased to 50 ms during the 75% validity condition. This is about the same magnitude than observed in human psychophysical experiments using the same stimuli and paradigm, after verbal instructions (Wegener et al. 2008), suggesting that the monkey quickly learned to use the cue information for allocating attention.

If reward amount is not considered, this significant, cue validity-dependent RT effect may seem surprising at first glance, because a high number of incorrectly cued changes is expected to promote distributed rather than spatially selective attention. Yet, non-binary rewarding allowed to select a 50% reward benefit for fast RTs and a 33% loss for slow RTs, compared to medium RTs. With decreasing cue validity, therefore, considering the cue information not only helps to more often get a high reward in a single trial, but at the same time, compensates a probable loss in average reward amount on the long run. In fact, in valid trials the monkey significantly increased its ratio of highly rewarded trials from 0.57 in the 90% validity condition to 0.74 in the 75% validity condition (*T*-test, *P* < 10^−3^, *N* = 6) (Fig. 3*D*). This was at the expense of reward in invalid trials, which showed an increase in the ratio of lowly rewarded trials from 0.07 to 0.3 (*P* < 10^−4^). This focusing on the cued targets allowed the monkey to even achieve a slight increase in average amount of reward per trial (*P* = 0.065), despite the reduction in cue validity (Fig. 3*E*).

### Example 4: Fine-shaping of behavior

Figure 4*A* introduces another change detection task requiring another monkey to select one out of more than 20 objects displayed in parallel (for simplicity only eight are shown in the Figure). The monkey was required to indicate a contrast change of the cued object, occurring at a random point in time between 0.7 and 4.7 sec after stimulus onset. It was trained using a non-binary reward scheme emphasizing fast RTs, similar to the approach described in the previous example. The monkey had just learned to deal with the high number of objects during the preceding training step, and therefore, the contrast change was kept easy to recognize. The next training step was to include changes at uncued objects - however, unlike the previous approach any uncued change would be followed by a change of the cued object later in the trial. As before, our question was whether rewarding of responses at uncued objects helps (or rather counteracts) training of the desired behavior, and furthermore, to what extent fine adjustments of reward amounts can be used to gradually shape behavior under this much more complex condition of visual stimulation.

**Figure 4.**
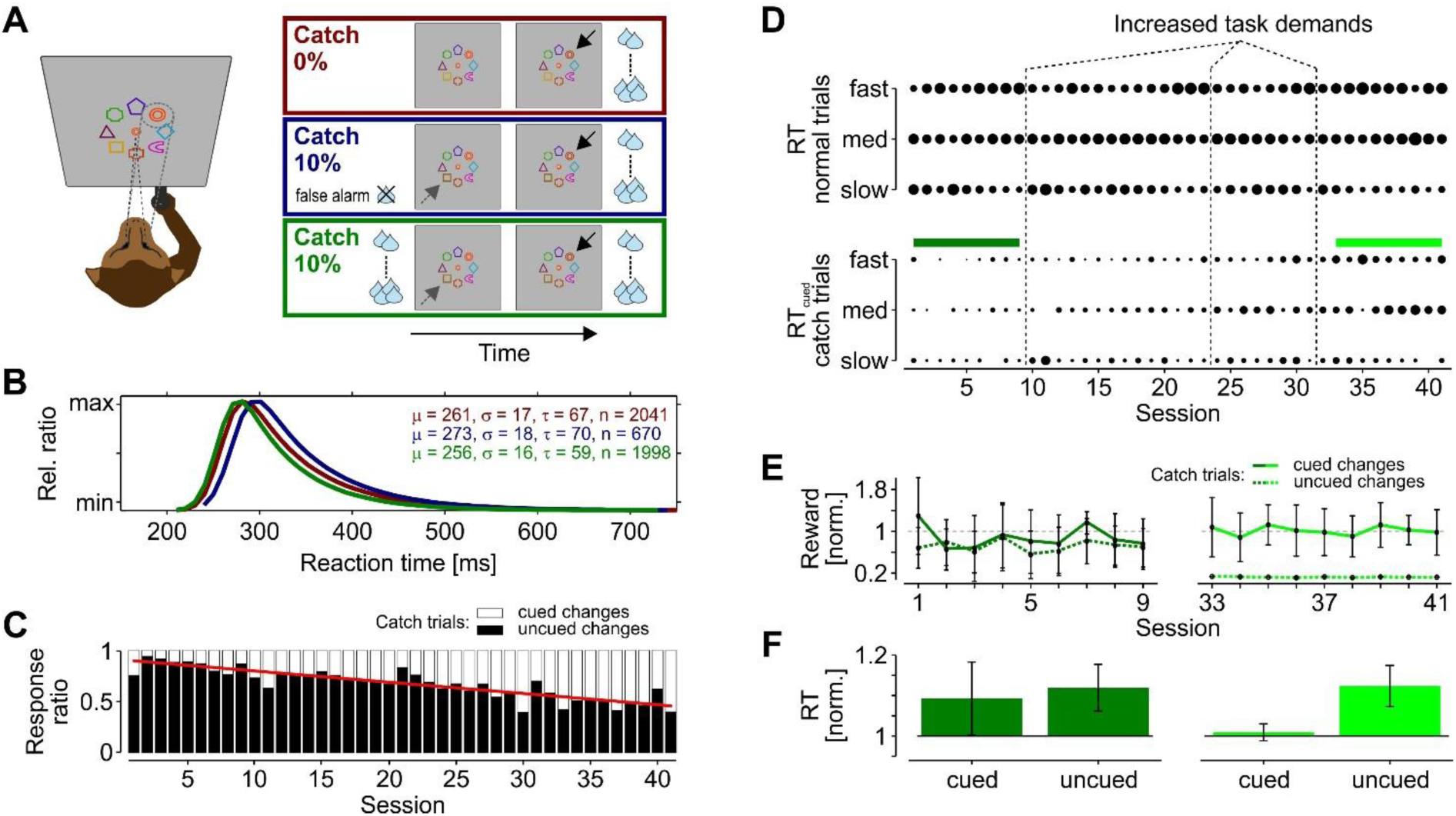
Reward-associated shaping of behavior. (*A*) Simplified visual display, showing eight out of more than twenty differently shaped and colored objects. The monkey’s task was to detect a luminance change at the cued object (displayed at the FP). In either 0% (red frame) or 10% of the trials, the luminance change at the cued object (straight black arrows) was preceded by a luminance change at an uncued object (gray dashed arrows). Detection of such uncued changes were treated either as false alarms and resulted in unrewarded trial termination (blue frame) or were rewarded as cued changes (green frame). Drop symbols indicate RT-dependent amount of reward. (*B*) Ex-Gaussian fits of RT distributions and corresponding fit parameters. Color code as in (*A*). (*C*) Ratio of responses to the cued and uncued change in catch trials during the course of 41 training sessions. Red line indicates a linear fit to the data. (*D*) Ratio of fast, medium, and slow responses to the cued change in normal trials (top) and in catch trials (bottom). Size of dots represents their relative number as compared to all correct responses in normal trials and in catch trials, respectively. Responses to uncued changes are considered for calculating catch trial ratios, but are not shown for simplicity. (*E*) Reward amount obtained in catch trials during sessions 1 to 9 (left, labelled with the same color as in (*A*, *B*, and *D*)) and during sessions 33 to 41 (right), after slowly increasing task demands and decreasing reward for responses to uncued changes. Reward amount is normalized to the reward amount obtained in normal trials during the same session. (*F*) Mean RT in response to cued and uncued changes in catch trials, normalized to mean RT during normal trials. Color coding as in (*E*). Error bars throughout figure, S.D.

We measured RTs during five sessions prior to the introduction of uncued changes. High, medium, and small rewards were provided for fast, medium, and slow RTs, respectively. Following these sessions, we introduced uncued changes in 10% of the trials (“catch trials”) and treated responses to these changes (UC responses) as false alarms, resulting in the termination of the trial without reward (as in the classic, binary approach). Ignoring these uncued changes and responding to the change of the cued object (C responses) later in the trial was rewarded as before. After two sessions with this feedback scheme, we switched to rewarding UC responses in the same way as C responses. The results of these two different approaches show two interesting features: First, in the two sessions during which UC responses resulted in the termination of the trial, the number of error trials (excluding fixation errors) increased by 50% as compared to the average performance in the five sessions without catch trials. Many of these errors were the consequence of a UC response, but even when disregarding catch trials, errors increased by 21%. Second, responses to correctly cued objects (disregarding catch trials) were slower than before. Fitting ex-Gaussians to the RT distributions before and after the introduction of catch trials revealed a rightward shift of the RT distribution by about 10 ms (Figure 4*B*, blue line). The data suggest that even with a catch-trial ratio of only 10%, the animal’s overall confidence with the task requirements was significantly attenuated when terminating trials after UC responses. In contrast, switching to rewarding UC responses helped to quickly re-establish the previous performance and the animal returned to about the same RT distribution as observed before the introduction of catch trials (Figure 4B, green line).

The key question is, however, whether this reward scheme supports the selective allocation of attention to the cued object (the desired behavior), or rather supports distributing attention over the entire stimulus array (the undesired behavior). With the exception of the very first training sessions introducing catch trials, a cued change occurred 750 ms after the uncued change at the earliest (cf. Material and Methods). For the monkey, focusing on the cued object increased the chance to achieve a fast response (i.e. getting a higher reward), but also bore the risk of a fixation error due to the longer trial time. Hence, because even slow UC responses would provide a safe reward, our reward scheme potentially facilitates the undesired behavior – a distributed rather than selective spatial attention. To obtain a detailed insight into the behavioral strategy of the animal, we introduced the following training steps rather slowly. For nine sessions, we left the task parameters unchanged and kept reward equal for UC and C responses. During the 22 sessions thereafter, we step-wise increased the task demands (reduction of contrast change, alignment of objects, double the number of catch trials) and reduced the amount of reward for UC responses, until finally UC responses received a very small reward only, independent of RT. Task demands and reward scheme were then kept unchanged for another nine sessions.

During the first sessions of this training period, in catch trials the monkey had a high ratio of UC responses (mean_S1-S9_: 82.25% ± 10.35). During this period, there was no strong need for the animal to allocate attention selectively to one stimulus, because the contrast change was relatively easy to recognize and responses to both cued and uncued changes often resulted in a high reward. Yet, with the adjustment of task demands and reward amounts there was an increasingly larger benefit for selectively attending to the cued stimulus, and the monkey started to respond more and more frequently to the cued change later in a catch trial (Fig. 4*C*). Comparing the first and the last nine sessions of the training, this change in behavior was highly significant (*T*-test: *P* < 10^−12^).

A detailed analysis of RTs and obtained reward provides a deeper insight into the factors likely responsible for this behavioral adaptation. Figure 4*D* shows the performance of the animal in each of the sessions, in terms of relative fractions of fast, medium, and slow C responses (resulting in high, medium, and low reward, respectively), separately for normal and catch trials. During the first nine training sessions, in normal trials the monkey responded more and more frequently with a fast RT, and reduced the number of trials with slow RT (Fig. 4*D*, upper panel). After increasing the task demands, each time performance dropped initially and then rose again during subsequent sessions, until the monkey eventually managed a high ratio of fast, highly rewarded responses with the final task demands.

Obviously, using the information provided by the cue got more and more beneficial to reach a high reward regime. This increased attention to the cued item is mirrored by the performance level in catch trials. At the beginning of the training, the monkey only rarely responded to the cued item in catch trials, and if so, managed only a small or medium reward in the majority of trials, suggesting a more distributed rather than selective attention. With increasing task demands, however, it responded more often to the contrast change at the cued object later in a catch trial, and frequently achieved a fast, highly rewarded RT (Fig. 4*D*, lower panel). From the perspective of the animal, this behavior was highly rational, as shown by the analysis of obtained reward (Fig. 4*E*). With low task demands and a more distributed attention, the monkey managed to get about equal amounts of reward for both UC and C responses in catch trials. Yet, as compared to normal trials the amount was significantly less after UC responses (72% of mean per-trial reward in normal trials, *T*-test: *P* < 10^−4^, *N* = 9) and slightly but not significantly less after C responses (89%, *P* = 0.512). In accordance with this, RTs to both cued and uncued changes were significantly slower in catch trials than in normal trials (*P* < 0.016 for both) (Fig. 4*F*, left panel), indicating that uncued events captured some attention, such that the monkey was generally slower during catch trials (even when disregarding the uncued change and responding to the cued item later in a trial). In contrast, at the end of the training, when task demands were higher and UC responses received a small to very small reward, the monkey strongly focused its attention on the cued item. In catch trials, it achieved as fast responses to the cued change as in normal trials (*P* = 0.229), and managed the same reward amount (*P* = 0.674) (Fig 4, *D* and *E*, right panels). UC responses were significantly slower (*P* < 10^−4^) than C responses, and were delayed by 36 ms on average. Thus, the animal learned to select one out of more than 20 displayed items and allocated spatial attention in the desired manner within a very acceptable time span.

Summarizing these four examples, NB-PRT proved to be effective for training both simple and complex task rules by emphasizing the positive behavior instead of relying on errors. The non-binary rewarding and the waiver of trial terminations for new task rules allows to keep the animal’s motivation on a high level and to effectively guide the animal towards the desired behaviors. Moreover, by segmenting the reward it provides both a more differentiated feedback to the animal and additional behavioral data to the trainer, and thus allows to adapt task demands or introduce new task rules at optimally effective points in time.

## Discussion

PRT is a widely used tool for training animals to cooperate for husbandry or scientific purposes. It might be used to train animals on entering a compartment or presenting an arm or leg for blood sampling, moderate aggressive or affiliative behavior amongst individuals, or as environmental enrichment (for review: Coleman and Maier 2010; Laule et al. 2003; Prescott and Buchanan-Smith 2003; Rogge et al. 2013; Schapiro et al. 2003; Westlund 2015). PRT has also been successfully used for automated procedures to assess cognitive abilities prior to training in a laboratory environment (Fagot and Paleressompoulle 2009), and for automated, voluntarily performed training procedures inside the animal facility (Calapai et al. 2016; Tulip et al. 2017).

Training animals on complex cognitive tasks for neuroscientific purposes faces some very specific additional challenges: first, animals are outside the familiar environment of their home cage or stable, in a laboratory environment; second, they need to be restrained by means of a primate chair and usually a head holder; third, the desired behavior relies on interpreting rather abstract sensory cues; and forth, they are required to perform many (usually several hundreds) consecutive trials to allow meaningful data collection. Given these particularities, successful training in reasonable time depends on generally accepted guidelines on the one hand, like rigid planning of training steps and prudent handling of the animal in the laboratory (Smith et al. 2006; Tardif et al. 2006). On the other, it requires specific approaches to assure the animal’s motivation and cooperation to perform the behavioral task it is thought to be trained on. PRT has been successfully used to train monkeys since the early years of neurophysiological recordings in awake, behaving animals (Evarts 1968; Goldberg and Wurtz 1972; Lemon and Porter 1976; Mountcastle et al. 1975; Wurtz 1969). Yet, with increasingly more sophisticated and demanding tasks to study higher cognitive functions and complex motor behaviors, training procedures get more difficult and time consuming, and may face unforeseen pitfalls potentially causing additional delays.

The method we here describe is a modification of the classic PRT approach used in the field. It was partly motivated by recent studies showing that monkeys integrate information about reward probabilities to bias their choices in free-choice paradigms (Feng et al. 2009; Kubanek and Snyder 2015; Rorie et al. 2010), and quickly learn selection rules based on stimulus-reward associations (Gaffan et al. 2002; Lennert and Martinez-Trujillo 2011). NB-PRT relies on this natural expertise and uses it for interacting with the animal. NB-PRT can be applied both for training a new task and for maintaining high performance levels when learning has finished. Based on the experience gathered, we identified three general hallmarks of this approach for laboratory training of both simple and complex cognitive tasks.

The first is that NB-PRT provides a more differentiated feedback to the monkey. During trial-and-error learning, monkeys learn how not to behave for preventing trial termination and rejection of reward. This learning of behavioral errors is critical for the progression of the training and its overall success (Sutton and Barto 1998). Yet, training of complex tasks involves introduction of new error sources at some critical steps. For the animal, such unfamiliar errors easily cause uncertainty with the current task rules, not only regarding the new task component but also with respect to already established behaviors (cf. Figure 4*B*). Association of specific error types with specific acoustic feedback signals may help to label error sources, but to our experience, there is not much reason to strongly rely on this. NB-PRT provides the opportunity to introduce such new error sources in the soft way and to keep the animal’s confidence with the general task rules. Instead of trial termination and rejection of reward, undesired behavior related to a new task rule is rewarded, but much lesser than the desired behavior (cf. Example 1 in Results). This makes the feedback more distinct: Previously learned error sources still cause trial termination, but the new error source is systematically ‘taught’ to the animal based on small rewards. After the animal learned the desired behavior (by figuring out how to get the most reward), the reward scheme can eventually be reset by fully rejecting reward for the undesired behavior first, and subsequently associating it with trial termination, such that new reward-behavior associations may be defined for the next training step.

The second hallmark is an increase in the learning rate and a more purposeful shaping of behavior. Because trial-and-error learning with binary rewarding cannot signal more than ‘right’ and ‘wrong’, unsuccessful trials will make the difference for the animal to adapt its behavior. Yet, their relative number must not exceed a critical ratio, to prevent a loss in task confidence and in the overall motivation for cooperation. This causes a conflict, because keeping their number within a critical range also limits the number of trials for learning. With a graded reward, it is possible to put emphasis on correctly performed trials. By a careful choice of criterions for e.g. high, medium, and low reward, NB-PRT provides the animal with a larger number of informative trials (Example 2), and guides it towards the desired behavior more purposefully than error-based learning (Examples 3 and 4).

The third hallmark is the introduction of a ‘gambling factor’ for tasks of otherwise uniform structure. As mentioned previously, in neuroscience research monkeys are usually required to perform hundreds of consecutive trials. Even when focusing on individuals that are very good performers in the laboratory, there is not much reason to believe that performing the same task again and again is constantly thrilling. NB-PRT provides the opportunity for the monkey to ‘win’ a trial by rewarding very good performance better than medium performance. By associating the task outcome with the animal’s performance it makes the reward more unpredictable and as such, provides a lasting incentive for the animal to stay focused even when learning has finished (see also Karolina Westlund’s commentary on variable-ratio schedules (Westlund 2012)).

Like any other approach, however, successful application of NB-PRT requires careful planning of the individual training steps, identification of possible pitfalls, and taking the perspective of the animal. When based on careful daily data inspection, it constitutes a powerful tool that provides more options to guide the animal’s behavior. Because it is based on success rather than failure, it also provides particular advantages for preventing animals to develop very low or very high error tolerance, and constitutes an important and promising approach for refining laboratory methods for non-human primates.

## Acknowledgements

Supported by Deutsche Forschungsgemeinschaft (DFG) grants WE 5469/2-1 and WE 5469/3-1, a ZF grant from the University of Bremen, and a scholarship from the German Academic Scholarship Foundation. The authors acknowledge support by Sally Lott, Miriam Schillner, Peter Bujotzek, Ramazani Hakizimani, and Katrin Thoß regarding various aspects of the study. We thank Eric Drebitz for comments on an earlier draft.

## Author contributions

D.W. designed research; B.F. and D.W. performed experiments; B.F. and D.W. analyzed data; B.F. and D.W. interpreted results of experiments; B.F. and D.W. prepared figures; D.W. wrote the article; B.F. and D.W. approved final version of the article.

